# Consensus and conflict among ecological forecasts of Zika virus outbreaks in the United States

**DOI:** 10.1101/138396

**Authors:** Colin J. Carlson, Eric Dougherty, Mike Boots, Wayne Getz, Sadie Ryan

## Abstract

Ecologists are increasingly involved in the pandemic prediction process. In the course of the Zika outbreak in the Americas, several ecological models were developed to forecast the potential global distribution of the disease. Conflicting results produced by alternative methods are unresolved, hindering the development of appropriate public health forecasts. We compare ecological niche models and experimentally-driven mechanistic forecasts for Zika transmission in the continental United States, a region of high model conflict. We use generic and uninformed stochastic county-level simulations to demonstrate the downstream epidemiological consequences of conflict among ecological models, and show how assumptions and parameterization in the ecological and epidemiological models propagate uncertainty and produce downstream model conflict. We conclude by proposing a basic consensus method that could resolve conflicting models of potential outbreak geography and seasonality. Our results illustrate the unacceptable and often undocumented margin of uncertainty that could emerge from using any one of these predictions without reservation or qualification. In the short term, ecologists face the task of developing better post hoc consensus that accurately forecasts spatial patterns of Zika virus outbreaks. Ultimately, methods are needed that bridge the gap between ecological and epidemiological approaches to predicting transmission and realistically capture both outbreak size and geography.

## Introduction

In the urgent setting of pandemic response, ecologists have begun to play an increasingly important role^1^. Ecological variables like temperature and precipitation often play just as important a role as socioeconomic risk factors in the vector-borne transmission cycle, governing key parameters including transmission rates, vector lifespan, and extrinsic incubation period^2^; the statistical relationships among these variables can be exploited to develop predictive frameworks for vector-borne disease outbreaks. These models are often developed for mechanistic prediction at local scales, but ecologists have recently begun to play a more important role in predicting the overall possible distribution of emerging infections. Ecological niche modeling is a typically phenomenological method that correlates occurrence data with environmental variables to make inferences about the geographic boundaries of potential transmission^3^. Within niche modeling approaches, there are conflicting views regarding which algorithms are appropriate to use in the context of particular applications^4-6^, and consensus methods have hardly advanced beyond basic model averaging^7^. As an increasingly popular alternative, mechanistic ecological models have been developed that extrapolate geographic projections from experimental results^8^, but these can be data-intensive and highly sensitive to parameterization^8-10^. In theory, the two approaches-phenomenological and top-down, or mechanistic and bottom-up-should be roughly congruent when implemented with sufficient data and predictors, as they approximate the same pattern^11^. Yet discrepancies between the two approaches in practice highlight a tension in species distribution modeling, between deductive approaches that infer ecology from observed broad-scale patterns, and inductive approaches that scale ecological experiments to predict real patterns. In the context of pandemic response, the trade-off has acute stakes: early access to ecological predictions can help pandemic efforts, but inaccurate information based on limited data could drive misallocation of public health resources.^12^. Thus, there is a clear need to develop better consensus methods, but even before that, a need to understand the epidemiological implications of the differences among model-building approaches.

The Zika virus (henceforth Zika) pandemic that was first detected in Brazil in 2015 highlights the unusual and sensitive challenges of pandemic response. A number of characteristics make Zika unique from a public health standpoint, including its rapid spread through the Americas after a slow, multi-decade spread from Africa through Asia; the appearance of a sexual route of transmission, a rare feature for a vector-borne pathogen; and perhaps most importantly, the appearance of high rates of microcephaly, and more broadly the emergence of Zika congenital syndrome. At least 11,000 confirmed cases of Zika have affected pregnant women, leading to roughly 10,000 cases of birth defects, including microcephaly,^13^. As of April 6, 2017, a total of 207,557 confirmed cases of autochthonous transmission (out of 762,036 including suspected cases) have been recorded in the Americas^14^. Moreover, Zika is exceptional among vector-borne diseases in that it has developed a sexual pathway of transmission in humans (comparable examples, such as canine leishmaniasis, are incredibly rare^15^). The rapid spread of Zika virus from Brazil throughout the Americas has posed a particular problem for ecologists involved in pandemic response, as several different ecological niche models (ENMs)^12, 16, 17^ and a handful of mechanistic forecasts^10^ have been developed to project the potential full spread of the pathogen. So far, autochthonous transmission has been recorded throughout most of Central America and the Caribbean, with cases as far north as the southern tips of Texas and Florida. The Zika virus (henceforth Zika) pandemic that was first detected in Brazil in 2015 highlights the unusual and sensitive challenges of pandemic response. A number of characteristics make Zika unique from a public health standpoint, including its rapid spread through the Americas after a slow, multi-decade spread from Africa through Asia; the appearance of a sexual route of transmission, a rare feature for a vector-borne pathogen; and perhaps most importantly, the appearance of high rates of microcephaly, and more broadly the emergence of Zika congenital syndrome. At least 11,000 confirmed cases of Zika have affected pregnant women, leading to roughly 10,000 cases of birth defects, including microcephaly,^13^. As of April 6, 2017, a total of 207,557 confirmed cases of autochthonous transmission (out of 762,036 including suspected cases) have been recorded in the Americas^14^. Moreover, Zika is exceptional among vector-borne diseases in that it has developed a sexual pathway of transmission in humans (comparable examples, such as canine leishmaniasis, are incredibly rare^15^). The rapid spread of Zika virus from Brazil throughout the Americas has posed a particular problem for ecologists involved in pandemic response, as several different ecological niche models (ENMs)^12, 16, 17^ and a handful of mechanistic forecasts^10^ have been developed to project the potential full spread of the pathogen. So far, autochthonous transmission has been recorded throughout most of Central America and the Caribbean, with cases as far north as the southern tips of Texas and Florida.

In this study, we focus on the United States as a test system for exploring conflict between different model predictions. A Brazil-scale outbreak of Zika in the United States could be devastating; one model for only six states (AL, FL, GA, LA, MS, TX) found that even with the lowest simulated attack rate, Zika outbreaks could be expected to cost the United States over $180 million, and estimates under worse scenarios exceed $1 billion^18^. Consequently, a high priority has been placed on developing accurate models that capture socioecological suitability for Zika outbreaks in the United States^19, 20^. However, we suggest that the lack of a consensus among different models of spatial risk renders the literature less credible or navigable to policymakers, as predictions under certain conditions span a range from 13 counties at risk^12^ to the entire United States (Figure 1)^10, 21^. At the time of writing, the majority of public health agencies in the United States were preparing for the apparent eventuality of Zika, based either on no prior geographic information, or basic data on the range of Aedes mosquitoes^22, 23^. Millions of dollars have already been invested in state- and city-level Zika preparation, even in areas without recently-recorded Aedes presence, and pesticide spraying for vector control has already had unanticipated consequences, including killing millions of honeybees^24^. Domestic efforts to prepare for Zika are not unreasonable in the absence of a consensus prediction about Zika's likely final range; the continued importation of new cases into every state in the U.S. likely amplifies the perceived threat of local outbreaks, especially given the pathway of sexual transmission (which could conceivably start stuttering chains^25^ outside regions of vector-borne transmission). However, an informed response to Zika in the United States requires both a greater consensus about at-risk areas, and a more precise understanding of the uncertainty contained in different ecological forecasts.

**Figure 1.**
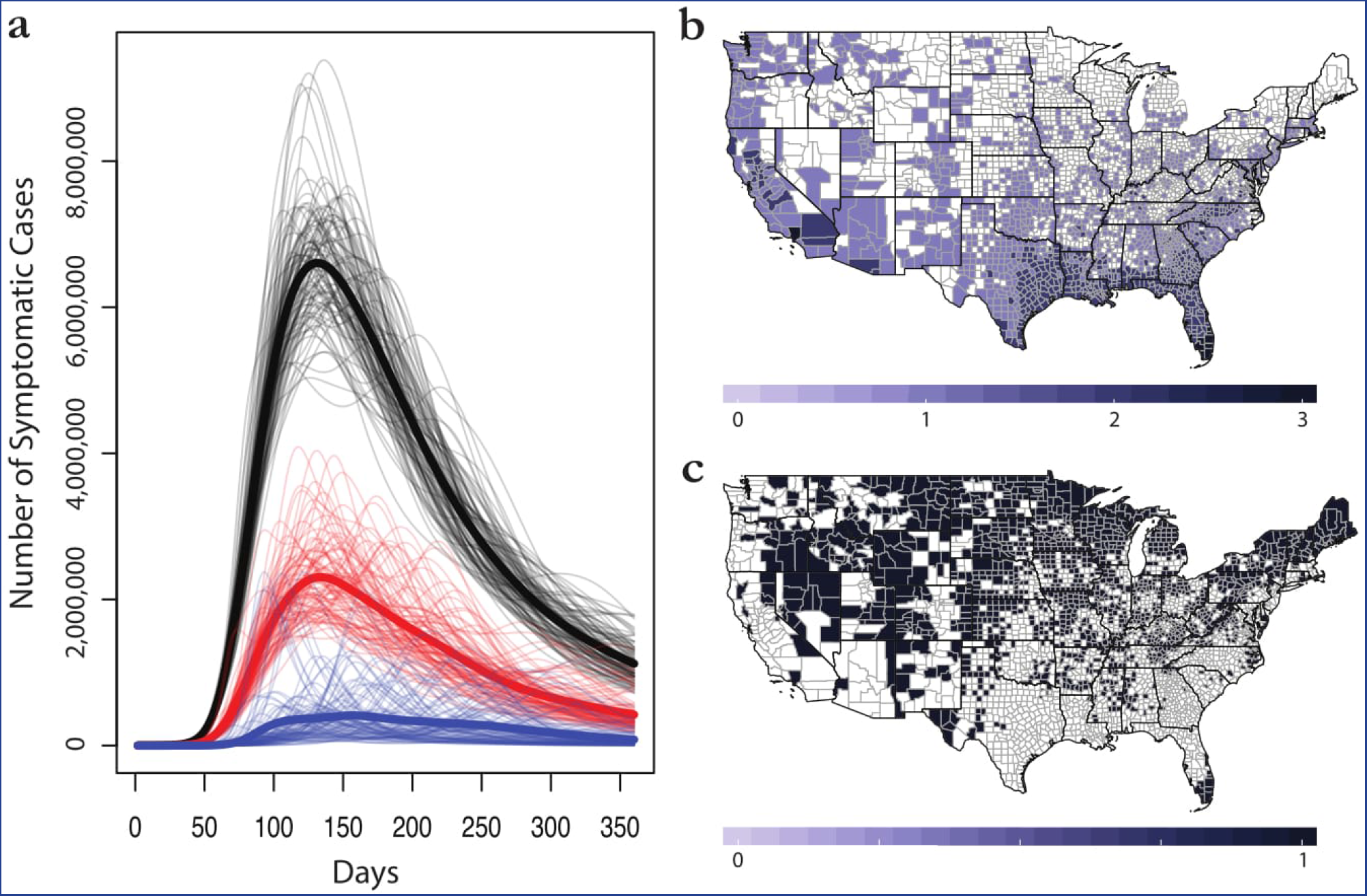
The margin of error in ecological niche models for Zika virus. (a) Average epidemiological forecasts associated with county data for Carlson (*blue*), Messina (*red*), and Samy (*black*), against a backdrop of overlapping individual simulations for each (grey). (b) The individual predictions of each model are given as presence or absence values; a maximum score of 3 indicates all models agree on presence, while a score of 0 indicates all models agree on absence. (c) Has consensus been achieved? At the county scale, dark blue indicates consensus among niche models; white indicates controversy. Maps were made in R 3.3.2^63^, using U.S. Census shapefiles.

Ecological niche models for vectors and pathogens are commonly used as an underlying foundation in epidemiological models, or more broadly, spatial studies in public health and policy work (including in the Zika literature^19, 26-28^) In this study, we highlight the unavoidable - but usually, unacknowledged - downstream consequences of model selection in those cases, and illustrate the lack of any one clear way to resolve conflict among published, peer-reviewed ecological studies. To expose this problem more clearly, we compare four published ecological predictions for the extent and duration of possible Zika virus transmission in the United States, and overlay generic epidemiological forecasts to measure the impacts of model differences. In doing so, we examine the scale of epidemiological uncertainty introduced at five scales:

1. Different environmental variable selections for a given niche modeling approach^17^
2. Differences among published ecological niche models
3. Differences between phenomenological^12, 16, 17^ and mechanistic^10^ approaches
4. Differences driven by parameterization of Bayesian mechanistic models^10^
5. Differences in how population-at-risk is aggregated from the niche models for epidemiological simulations

In the process, our exercise shows that relying on any one ecological model adds a hidden layer of uncertainty to epidemiological forecasts, indicating the need to develop better consensus methods—and to develop ecological and epidemiological tools in a more integrated approach that better approximates observed outbreak patterns.

## Results

Ecological forecasts for Zika suitability span the range of thirteen counties to all 3108 counties in the continental United States (Table 1), and this uncertainty (unsurprisingly) produces tremendous downstream variation in outbreak size. For ENM-based projections, the margin of error among mean trajectories spans an order of magnitude, with a total difference of 168 million cases between Carlson and Samy (Figure 1). Areas predicted by other methods to be at the greatest risk from Zika virus are roughly agreed upon among the models, with southern California and the Gulf Coast represented most significantly as outbreak hotspots among the three models. Agreement among all three models is limited, but is most significantly clustered in these areas, especially in the southern tip of Florida and Los Angeles County. While we assumed that aggregating risk at the county level could potentially absorb some of the spatial uncertainty of models and decrease differences between them, we found that it actually substantially exacerbated the observed differences among them (Table 2). This was perhaps most notable in the most populous counties, such as Los Angeles county (see supplementary Figures S1-S3).

**Table 1.** A comparison of the different ecological forecasts. Four different methods, each performing well based on sufficient data and predictors, produce highly contrasting results. Out of a total of 3108 counties in the continental U.S., only five have experienced outbreaks (Cameron County, TX with 6 cases of local transmission in 2016; Miami-Dade, FL with 241; Palm Beach, FL with 8; Broward County, FL with 5; and Pinellas County, FL with 1)^33, 62^. Accuracy values were calculated from the confusion matrix of observed outbreaks against predicted suitability. The Carlson model comes closest to predicting the geography of those outbreaks most accurately; but all epidemiological models “overpredict” the number of suitable counties based on the current extent of outbreaks. (Mordecai results are split for the highest bound with minimum temperatures, and the lowest bound for maximum temperatures, to give the full range of predictions. Self reported AUC values are shown not as a comparative measure of accuracy, but simply as the self-reported accuracy of the studies. Samy *et al*. used the Partial ROC in place of the AUC but did not report values. NA = Not Applicable; NR = Not Reported)

|  | Carlson | Messina | Samy | Mordecai (97.5% min) | Mordecai (2.5% max) |
| --- | --- | --- | --- | --- | --- |
| $n_{points}$ | 242 | 323 | 168 | NA | NA |
| $n_{predictors}$ | 15 | 6 | 15 | NA | NA |
| AUC | 0.970 | 0.829 | NA | NA | NA |
| Counties Predicted | 13 | 465 | 1616 | 1937 | 3108 |
| Accuracy | 99.6% | 85.2% | 48.2% | 37.8% | 0.2% |
| County Population at Risk | 19,653,445 | 95,359,408 | 270,249,781 | 218,444,263 | 320,957,062 |
| Mean Outbreak Size | 12,871,005 | 63,622,367 | 181,290,371 | 37,598,099 | 198,910,979 |
| Median Outbreak Size | 14,552,250 | 64,038,273 | 181,732,629 | 37,312,233 | 197,731,918 |

**Table 2.** Aggregating risk to the county scale can absorb some of the inherent spatial uncertainty of ecological niche modeling, but is itself an assumption that changes downstream impacts on the scale of outbreaks, as well as the scale of disagreement between models.

|  | Carlson | Partial | Messina | Partial | Samy | Partial |
| --- | --- | --- | --- | --- | --- | --- |
| Counties Predicted | 13 | 13 | 465 | 465 | 1616 | 1616 |
| County Population at Risk | 19,653,445 | 6,264,516 | 95,359,408 | 38,085,602 | 270,249,781 | 99,324,226 |
| Mean Outbreak Size | 12,871,005 | 4,195,326 | 63,622,367 | 25,897,671 | 181,290,371 | 66,345,567 |
| Median Outbreak Size | 14,552,250 | 4,262,636 | 64,038,273 | 26,307,445 | 181,732,629 | 66,850,610 |

Model parameterization has a considerable impact on downstream epidemiological results. The four models proposed by Samy, each with slightly different environmental variable selection (see the Methods), produced correspondingly different results (Table 3, Figure 2). Perhaps counter to our a *priori* expectations, adding more predictors produced broader projections and larger epidemics (not tighter-fit models); the model with all predictors (model 4) produced the largest epidemic, while the one with only climatic covariates (model 1) was in fact somewhat smaller than the Messina outbreak simulation. Model 2 (only social predictors) was only slightly more severe than model 4 (all predictors, which we use as the "Samy model" in all other cases). Adding more predictors increased the projected impacts most noticeably in Arizona and New Mexico; the projections in model 1, the most conservative, were very similar geographically to the Messina model, but with much more substantial range in the Pacific northwest.

**Figure 2.**
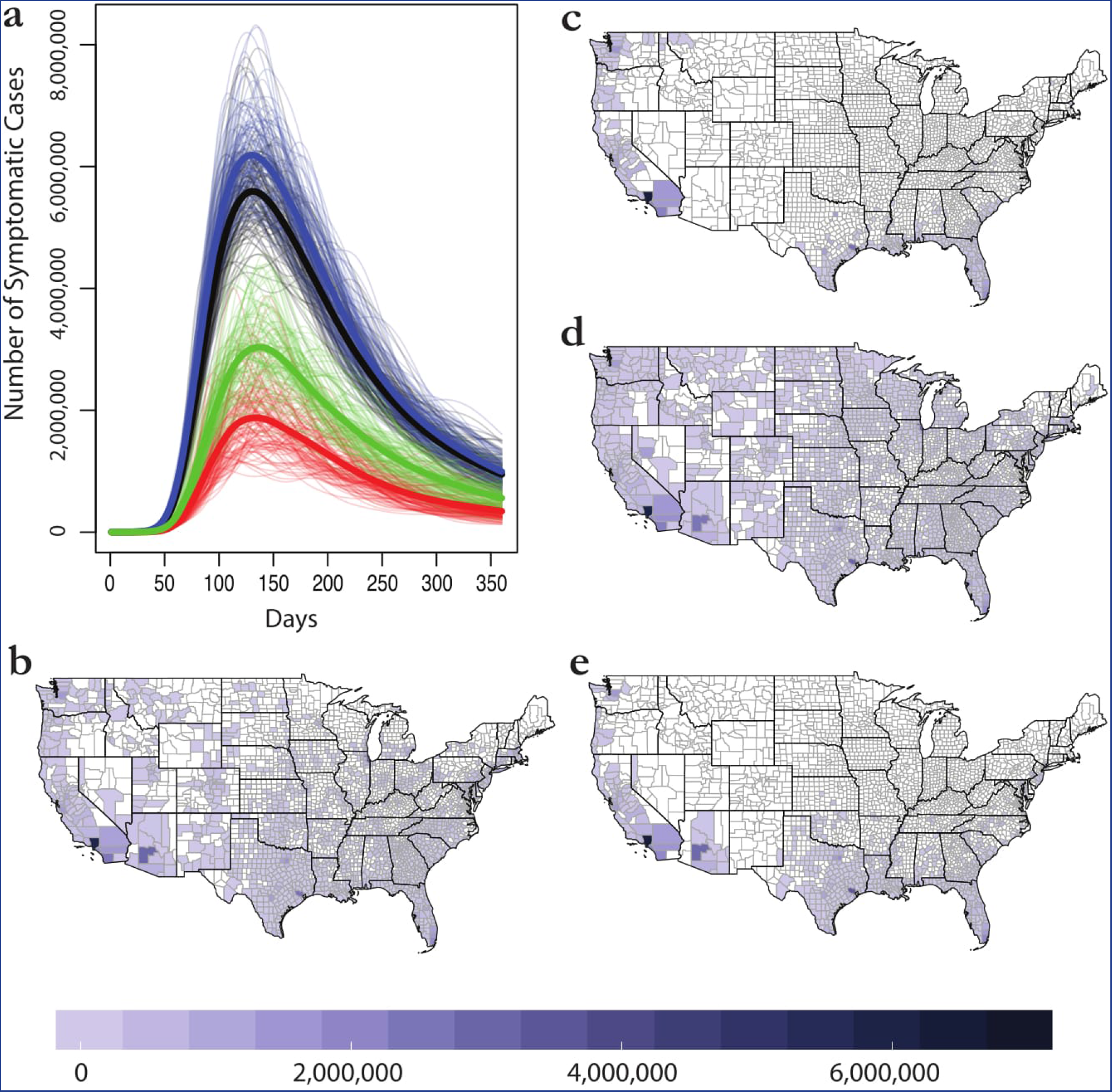
Variation within the Samy models. Outbreak trajectories are shown in (a) for models 1 (red), 2 (blue), 3 (green), and 4 (black). Bolded lines are mean trajectories. Final average case totals are then mapped for model 4 (b), the main model we discuss in the text and use in other comparisons, as well as models 1 (c), 2 (d), and 3 (e).

**Table 3.** Outbreak simulations exhibit greater than threefold variation in predictions among the four models presented in Samy.

|  | Model 1 | Model 2 | Model 3 | Model 4 |
| --- | --- | --- | --- | --- |
| Counties Predicted | 338 | 2197 | 670 | 1616 |
| County Population at Risk | 91,174,791 | 296,007,551 | 148,700,587 | 270,249,781 |
| Mean Outbreak Size | 59,561,603 | 197,553,462 | 98,314,605 | 181,290,371 |
| Median Outbreak Size | 59,561,222 | 197,759,983 | 98,770,516 | 181,732,629 |

**Table 4.** Majority rule based consensus models, meant to resolve uncertainty between the forecasts and provide a middle scenario. The main majority rule model combines the Carlson, Messina, and Samy forecasts; the seasonal majority rule model assigns monthly suitability values to that forecast, based on the minimum temperature 97.5% Mordecai model.

|  | Majority Rule | Seasonal Majority Rule |
| --- | --- | --- |
| Counties Predicted | 383 | 370 |
| County Population at Risk | 93,195,970 | 87,632,865 |
| Mean Outbreak Size | 60,259,904 | 24,267,441 |
| Median Outbreak Size | 60,940,111 | 24,407,885 |

A roughly comparable range of predictions to the span of the Carlson, Messina, and Samy models is contained within the entire span of possible implementations of Mordecai's Bayesian model (Figure 3). Whereas ENM approaches indicate a somewhat restricted geographic range for possible outbreaks, the Mordecai model suggests that even in a conservative scenario (using minimum temperatures, and 97.5% posterior probability), the majority of *Aedes aegypti'*s range is at least seasonally suitable for Zika transmission. A far greater range of variation is contained within the minimum-temperature-based model scenario, which encompasses roughly half of the land area of the continental U.S. In contrast, the three scenarios based on maximum temperatures are geographically indistinguishable, though worsening projections do extend the seasonality of transmission and thereby produce somewhat longer-tailed epidemics (Figure 3a).

**Figure 3.**
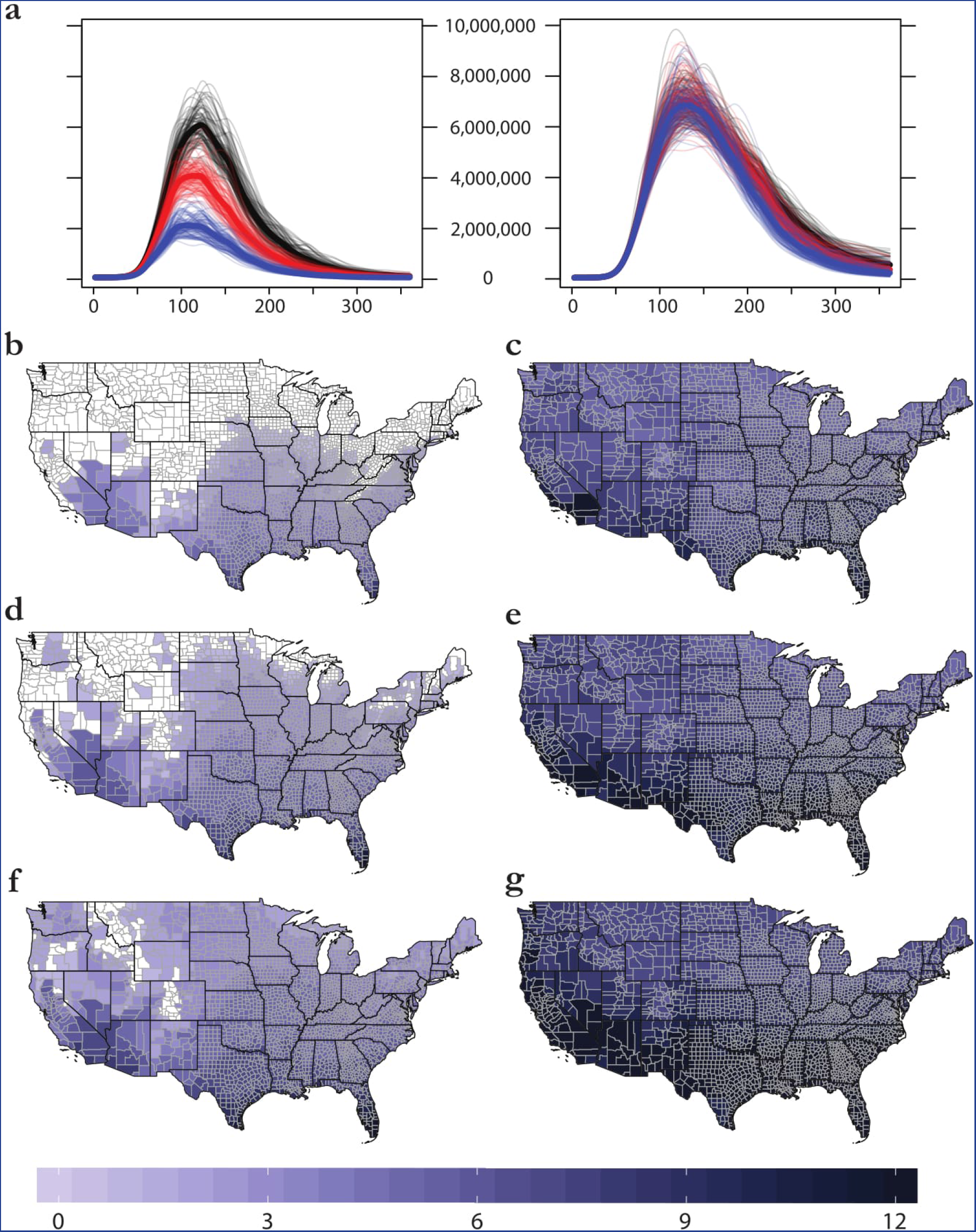
The margin of error within a single Bayesian mechanistic model for Zika virus, applied to minimum (left) and maximum (right) monthly temperatures. (a) 100 outbreak simulations for 97.5% (blue), 50% (red), and 2.5% (black) confidence intervals. (b-f) The number of months each county is predicted to be suitable for Zika virus transmission (R_0_ > 0) for 97.5% (b,c), 50% (d,e), and 2.5% (f,g) scenarios. Maps were made in R 3.3.2^63^, using U.S. Census shapefiles.

Among nine ecological scenarios considered (three niche models and six mechanistic scenarios), an overwhelming spread of possible epidemics could be predicted for the United States (Figure 4). The accompanying spatial pattern of case burden also varies between interpretations; while the spatial patterns are roughly identical for Carlson, Messina, and Samy, the temporal dimension introduced by mapping the Mordecai model onto monthly temperature grids dramatically affects how cases are ultimately distributed—and produces a reduction in epidemic size in some scenarios (Figure 5). In fact, the most conservative Mordecai scenario (97.5% confidence with minimum temperature) falls between Carlson and Messina in terms of case burden, despite predicting more than four times as many counties with transmission suitability as the Messina model. Across all models, forecasts predict that the majority of the case burden will still be seen along the Gulf Coast and in southern California. The advantage of interfacing ecology and epidemiology is especially evident here; for example, while the original Carlson *et al*. study noted the most significant suitable area was in southern Florida and failed to comment on the potential importance of southern California, the most significant epidemic predicted by most models is in Los Angeles county. The exception is the most conservative Mordecai scenario (Figure 2b), the only parameterization of that model in which Los Angeles is designated unsuitable for transmission—a fairly important discrepancy, given that the county is the most populous in the United States, and correspondingly contributes substantially to epidemic size in every other scenario (Figure 5a-c).

**Figure 4.**
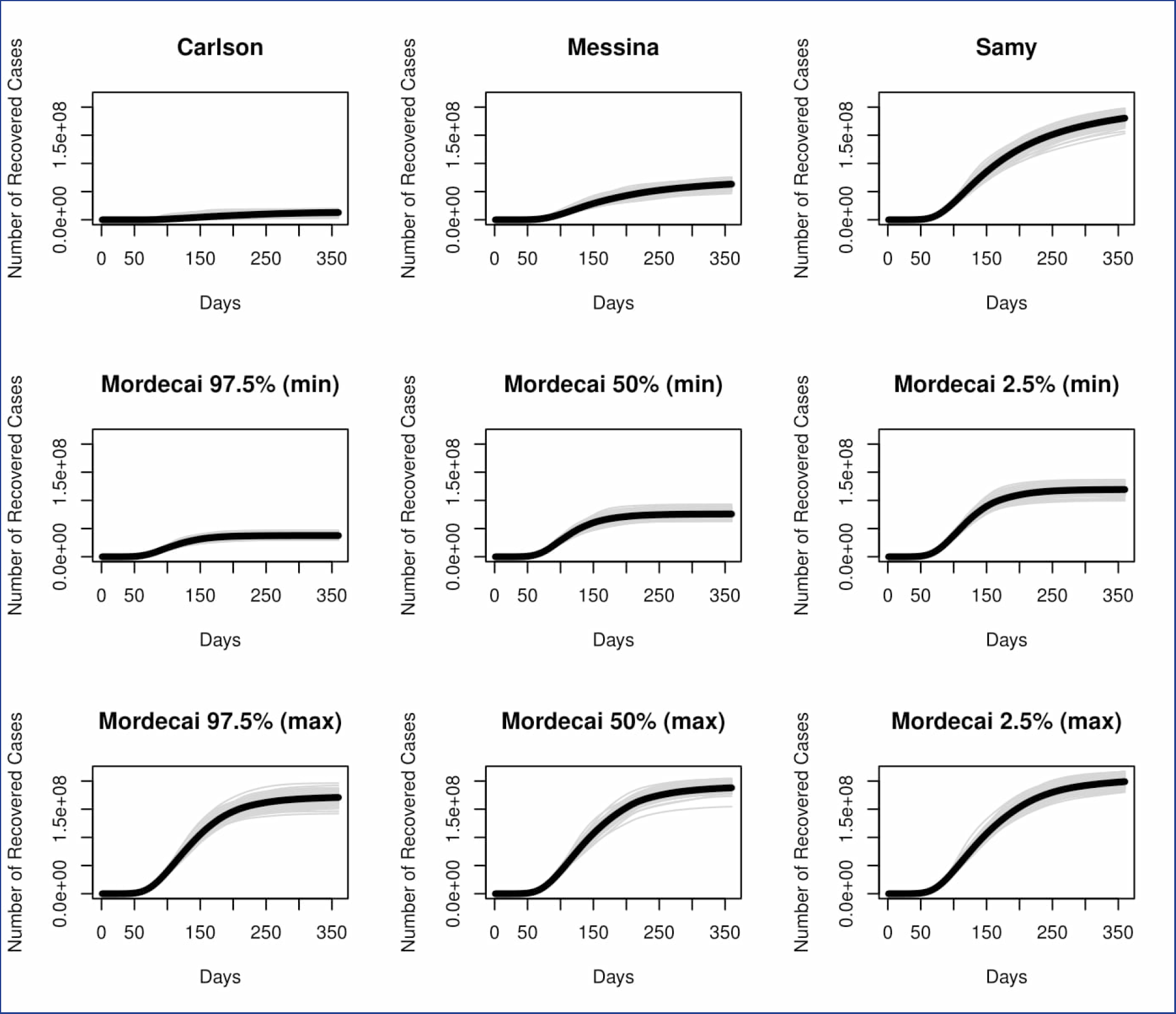
Nine possible trajectories for outbreaks in the United States: three based on ecological niche models, and six based on Bayesian mechanistic forecasts. (*y-axis on log scale*)

**Figure 5.**
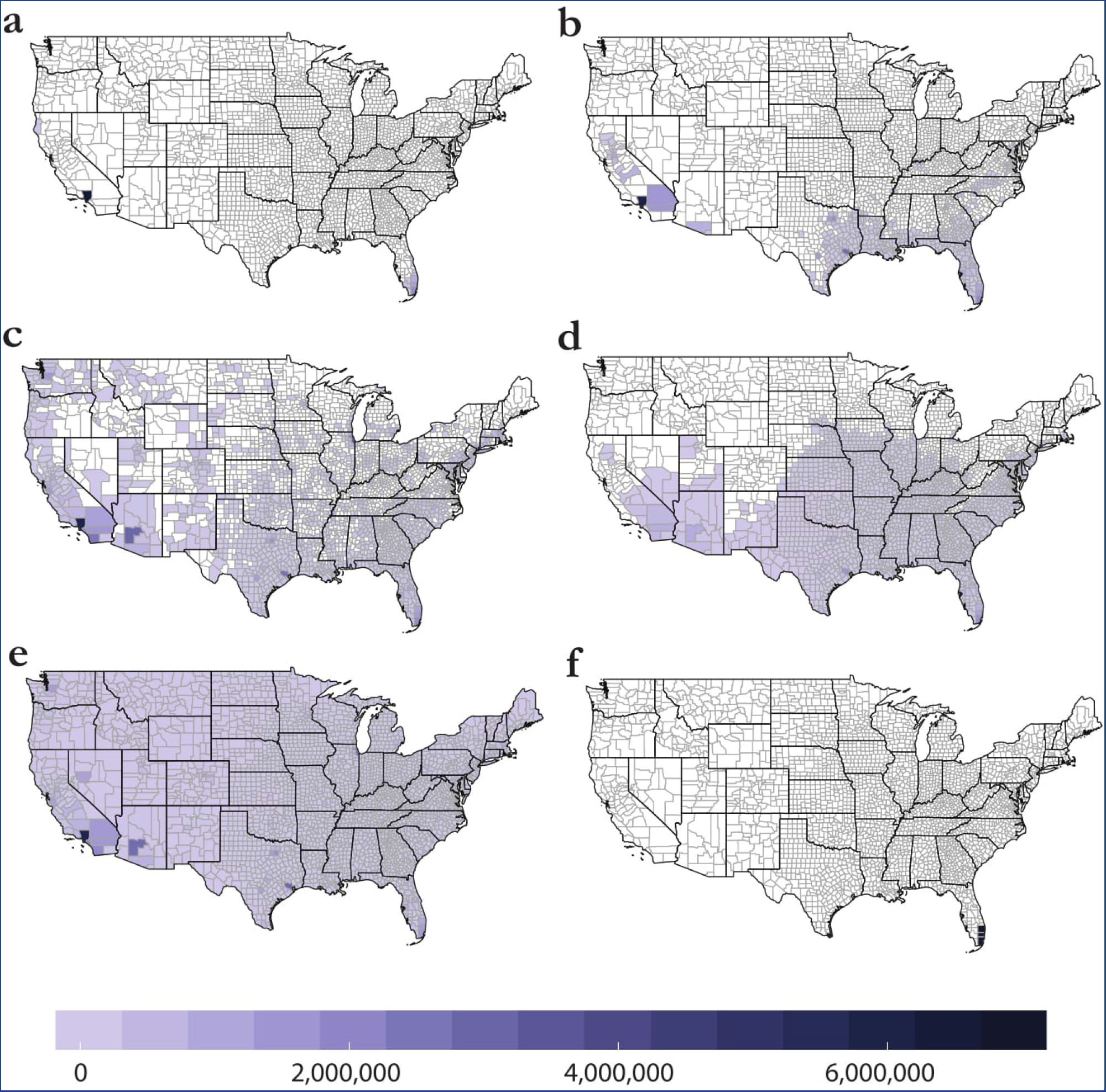
Case totals by county for (a) Carlson, (b) Messina, (c) Samy, (d), Mordecai 97.5% confidence (minimum temperatures), and (e) Mordecai 2.5% confidence (max temperatures), compared against (f) counties with reported autochthonous transmission in 2016 (three in Florida, one in Texas). Maps were made in R 3.3.2^63^, using U.S. Census shapefiles.

In an effort to illustrate a method of resolving these conflicting predictions, we present a final “consensus model” that incorporates all four modeling studies. Consensus methods are limited for ecological niche models^7^, so we adopt one possible approach: a majority rule at the county scale across Carlson, Messina, and Samy (i.e., in Figure 1b, any county value at or above 2 is “suitable,” and any below is “unsuitable”). Building on this “majority rule model,” for counties that are marked suitable by the ecological niche models, we superimpose the monthly transmission values from Mordecai's most conservative scenario, which most closely matches the geographic extent predicted by the ecological niche models (Figure 3a versus Figure 5a-c). This filtered “seasonal majority rule” algorithm incorporates the temporal dimension of transmission that is added by our implementation of the Mordecai model while maintaining consensus among the niche models.

The seasonal majority rule model produces a somewhat unsurprising pattern where year-round transmission is most common in the tropics, with seasonal transmission most important in the southeast United States, southeast Brazil, southeast China, and the Himalayas. Unsurprisingly, this produces a comparatively conservative outbreak prediction (Figure 7). The inclusion of the temporal component from the mechanistic model reduces case burden by almost two-thirds (Table 2), and excludes a handful of counties in the process (which were suitable in the ENM approach but not suitable for a single month in the mechanistic model). Most notably, Los Angeles county (which is suitable for no months of the year in the conservative Mordecai model) is excluded despite being suitable in all three ENMs, which contributes substantially to the overall reduction of projected case totals in the seasonal majority rule approach.

**Figure 6.**
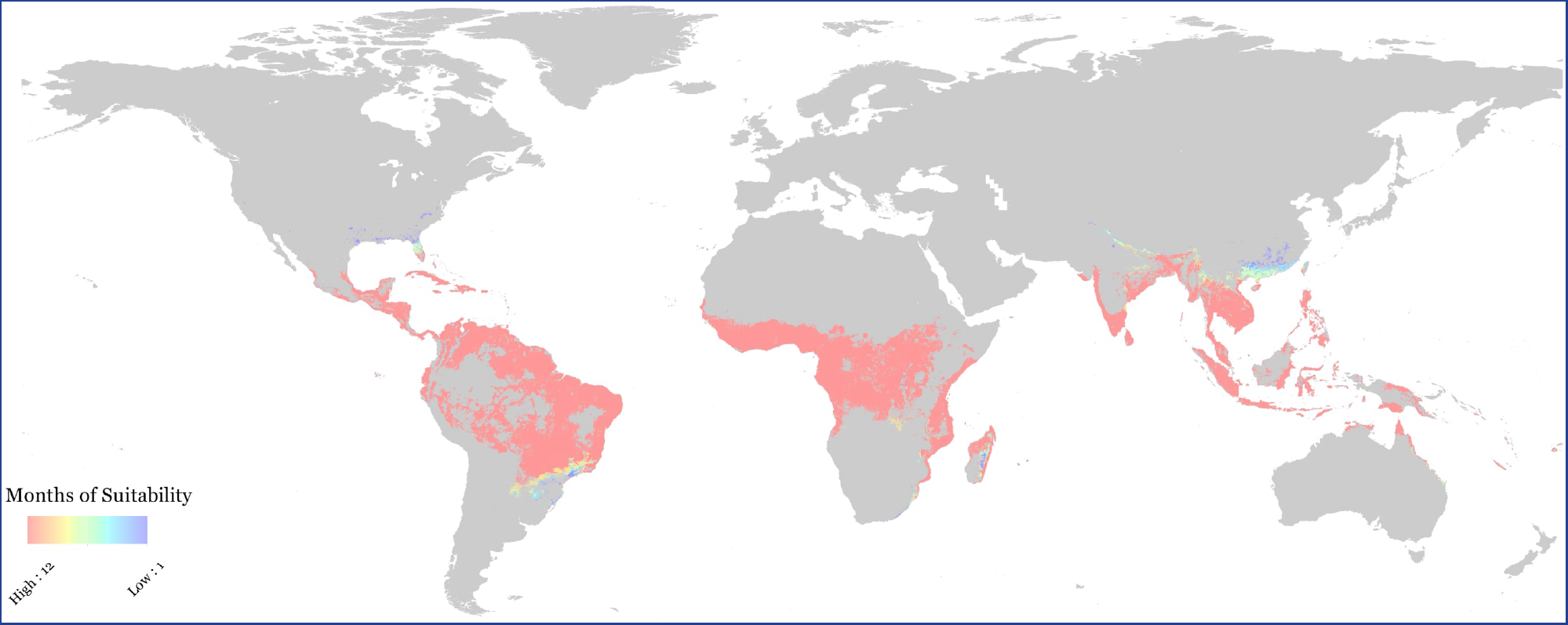
A global, consensus-based, seasonal (monthly) majority rule map of suitability for Zika virus transmission.

**Figure 7.**
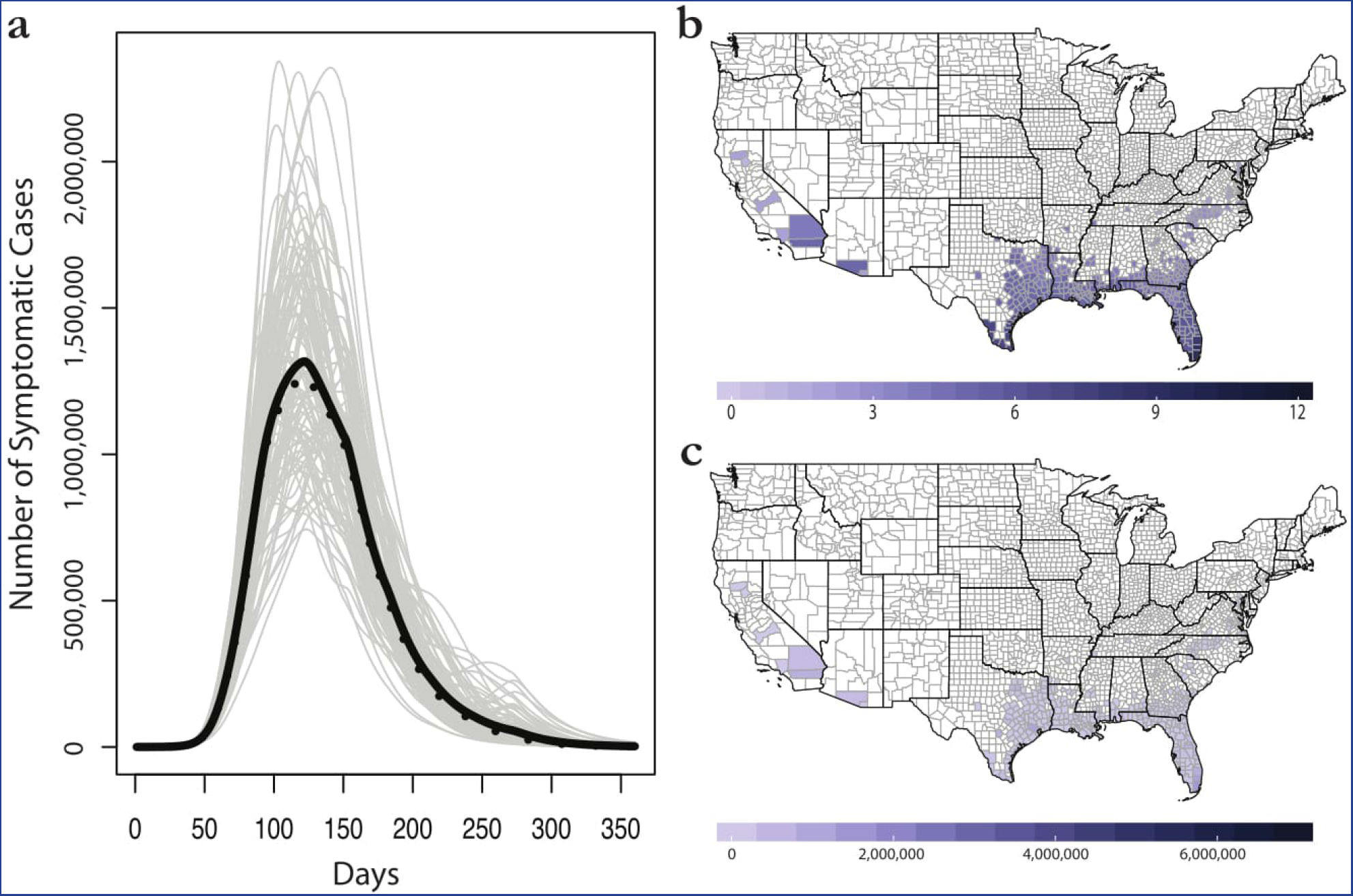
The seasonal majority rule method for consensus building across ecological forecasts. (a) Mean (black) and median (dashed) trajectories for 100 epidemic simulations. (b) The majority rule map: shading represents the number of months each county is marked suitable for outbreaks. (c) Final average case totals in the seasonal majority rule method. Maps were made in R 3.3.2^63^, using U.S. Census shapefiles.

## Discussion

### Clear Problems, No Easy Answers

By constructing epidemiological simulations on top of ecological niche models, we found that subtle differences among—and within—rigorous modeling frameworks can introduce substantial downstream variability in outbreak sizes and durations. Conflict among different published studies is the most immediately apparent problem, especially given the lack of a more sophisticated method of resolving these differences beyond the majority rule approach we use. However, for any given study, we show that internal model assumptions carry a level of uncertainty that is hard to understand just from looking at a “final model,” a fact that is readily apparent by comparing the different Samy models and Mordecai model parameterizations. (This can be a problem even in cases where no conflict exists among different published models; for example, one model of *Aedes aegypti* and A*e*. *albopictus* is most commonly used across purposes^29^, including frequently as an outer bound in epidemiological models.) Subjective model design issues like occurrence data collection and thinning, environmental variable selection, pseudoabsence generation, model algorithms, and threshold selection all introduce subjectivity into niche modeling that goes beyond basic issues of accuracy and exposes deeper strategic tensions in modeling (e.g., Levins' proposed tradeoff in modeling among realism, precision, and generality^30^). Mechanistic models are often designed as a response to that subjectivity but, as we highlight here, they also produce another conflicting result or set of results; moreover, Bayesian model parameterization still introduces downstream variability, possibly even more so than niche modeling.

We also found that the mechanistic models we examined produced much more inclusive predictions than any other model we considered (in accordance with work in parallel fields similarly suggesting mechanistic models favor generality and realism over precision^31^). To some degree, this conflict may expose an underlying tension between two different intentions of disease mapping. One paradigm focuses on accuracy (especially specificity), and follows a similar paradigm to mainstream invasion biology research in that it attempts to most accurately project the final boundaries of incipient range expansion. Overprediction and underprediction are weighted as equal problems in this approach; the task of appropriate allocation of clinical resources is equally impeded by both margins of error. An alternative paradigm assumes that a Type II error (excluding regions at risk of outbreaks) is of far greater significance than a Type I error (predicting risk for areas that remain unaffected), from a preparedness standpoint; and reacts especially to the stakes of under-prediction by targeting predictions at any area that could, theoretically, sustain outbreaks. In reality, all disease distribution models fall somewhere on a continuum between the two paradigms, and modelers following best practices are likely to produce primarily objective results. But to the degree that no forecasting effort is fully unsupervised, and basic decisions (like including or excluding current outbreak data) introduce opportunities for subjectivity, conflict between these two approaches is likely to be an ongoing disciplinary problem beyond Zika.

We note also that deliberate choices we made in the epidemiological models we included similarly produced a specific, and extreme, result. By setting mosquito populations high and using wide stochastic priors rather than tailored Zika outbreak parameters, we simulated unrealistic outbreaks on a scale even greater than seen in Brazil. (However, we note that a landmark study just published estimates that 12.3 million cases of Zika are expected every year in Latin America and the Caribbean; and while the United States was unassessed, northern Mexico was identified as a region of high variability and therefore high epidemic potential.^32^) Our methods here are meant to illustrate the full potential of hidden uncertainty that epidemiological models might inherit from ecological assumptions. The range of projected epidemics varies among the ENM approaches by more than an order of magnitude, but even the smallest outbreak prediction is still five orders of magnitude higher than the case totals observed during the last outbreak season (223 real cases versus roughly 12 million simulated cases). In practice, epidemiological models fitted to data may absorb some of the uncertainty between different ecological forecasts if outbreaks are constrained in areas of disagreement by additional socioecological factors. The scale of the problem is difficult to evaluate except on a case by case basis; at a minimum, we conclude that understanding the epidemiological implications of ecological uncertainty is a key step towards improving ecologists' performance in pandemic preparedness.

Ecological niche modeling is a comparatively new statistical method in ecology, and it has only recently been applied to emerging infectious diseases. In under two decades, the statistical power of ENMs has grown exponentially, especially as increasingly complex methods for machine learning have been applied to the problem. The dozen or so methods currently employed offer a wide palette of options for potential modelers to choose from, and compounded with the wide range of potential environmental and social covariates, seemingly limitless combinations of possible models can be produced from a single dataset, each of which is statistically rigorous enough to be published. Although guidelines exist for method selection and model tuning (e.g., variable selection), tremendous user-end creativity is still possible. High-profile targets, such as vector-borne and other zoonotic diseases, frequently inspire conflicting models, but in mainstream species distribution modeling research, the impacts of those conflicts are often treated in as an academic problem. For infectious disease mapping, such conflict has conspicuous stakes that produce downstream uncertainty for stakeholders, clinicians, and policymakers.

### Future Directions for the United States

Despite the disagreement between different modeling approaches and results, southern Florida and southern Texas clearly emerge across studies as the most at-risk regions of the continental United States for Zika virus outbreaks. This appears concordant with the broader consensus in public health research, especially given that these are already the only regions with a recent history of dengue outbreaks in the continental U.S.^33^ We also note that, in many of the models we considered, Los Angeles county emerged as a potential area of significant concern, especially given its dense population. But for the rest of the country, model disagreement is high and unresolved.

Given the wide suitable area suggested by the majority of models, the low totals of autochthonous cases in the continental United States still seems surprising. Epidemiological work supports the idea that the 2016 outbreak was not anomalously small; recent work estimated the *R*_0_ of the Miami-Dade outbreak in the low range of 0.5 to 0.8, and found that multiple introductions (an estimated 4 to 40) were a necessary precursor for an outbreak on the scale of the 256 cases in 2016^33^. Continued or larger outbreaks could be possible in the future if the high force of infection from traveler cases—which have so far been an order of magnitude more common in the U.S.—drives more significant outbreaks than the 2016 outbreak in Florida. More realistically, a number of factors likely prevent the United States from experiencing an outbreak on the scale that Brazil or Colombia experienced. Some are ecological; vector populations may be more strongly seasonal at higher latitudes, or the sylvatic cycle of Zika may be different in different parts of the Americas. The role non-human primates play in the transmission of Zika is still poorly understood^34, 35^, but the absence of monkey hosts could plausibly limit transmission in the United States. Lessons from chikungunya suggest that attention may need to be paid to potential alternate, sylvatic vectors and associated hosts^36, 37^, especially given the significant number of vectors that may be competent for Zika transmission in the United States^21^.

Other potential explanations for the limited spread of Zika through the United States are more social or socioecological in nature. In developed countries, household exposure is often secondary to outdoor exposure for *Aedes*, and in Miami-Dade county, it has been suggested that heterogeneity in outdoor exposure could have produced a much smaller, faster epidemic^38^. Other plausible explanations include better access to health care, preemptive vector control as part of Zika preparedness efforts, and significant fine-scale heterogeneity limiting mosquito populations in well populated areas (a factor that some models can accommodate^39^, but niche models at the global scale do not). The last of these is most easily addressed through ecological tools, and finer-scale validation of downscaled ecological models is an important next step for ongoing forecasting. At the county scale, more detailed GIS data are needed to identify probable areas of suitable vector density; identifying those areas can reduce the population at risk (used to parameterize models) from the population of an entire county down to just those living in high-risk (or non-zero risk) areas.

### Future Directions for Model Development

At the present time, the most common practice to address the ecology-epidemiology interface in the niche modeling literature is the use of population-at-risk (PAR) methods. Basic area-under-the-model population estimates are perhaps the simplest and most readily comparable possible epidemiological metric; only Messina *et al*. present a global PAR (2.17 billion people) based on their Zika virus niche model. Bogoch *et al*. revised that figure in a more regional assessment for Africa, Asia, and the Pacific that included traveler populations and a seasonal component to transmission, but to do so, substituted existing dengue models in place of actual Zika models.^26^ If implemented more frequently, population-at-risk methods could be a simple *post hoc* way of comparing different ecological forecasts. However, these methods might accidentally introduce more alarm than they communicate risk (just as using susceptible populations, without any associated model of transmission, is a fairly uninformative proxy for an epidemic projection in mainstream epidemiology). The exercise carried out here illuminates one of the primary weaknesses of ecological niche modeling methods; namely, though ENMs have great value for defining the plausible outer bounds of transmission, they are largely unable to clarify the distribution of risk within these suitable areas (except in rare cases where extremely specific populations at risk can be measured, e.g., rural poor livestock keepers at risk of anthrax^40^). Modeling approaches that more directly interface ecological and epidemiological concepts of risk and hazard are perhaps the “Holy Grail” of work at this interface, and approaches along these lines have recently been tested for hemorrhagic viruses in Africa^41, 42^. But we show here that uncertainty and subjectivity on the ecological side are propagated through approaches like these, with no clear solution.

The uncertainty at this interface represents a major deficiency in our ability to forecast disease spread. However, there are a number of potential of avenues of exploration that may help improve efforts to directly link epidemiological forecasts and ecological projections. On the epidemiological side of the problem, travel-based models have shown promise for other diseases^43^, and have been applied in a limited capacity with dengue models to predict Zika risk^26^. These types of models can be applied with Zika-specific niche models for more detailed forecasts of traveler-driven outbreaks at the edges of suitability. But a more detailed epidemiological link is needed between traveler force of infection and the scale of subsequent local outbreaks; so far, that causation has only been investigated in reverse.^44^ The role of sexual transmission also requires deeper investigation. Early work suggested sexual transmission might be a substantial factor explaining the explosive South American outbreak^45, 46^, but recent work has suggested sexually-transmitted outbreaks are unlikely^47^, even if sexual transmission increases the severity of vector-borne outbreaks^48^; others still argue these risks are “understated.”^49, 50^ Some work at the county level has already begun predicting Zika risk based on other sexually transmitted diseases^51^, but for this to be useful to policymakers, a basic and accurate model of importance of sexual transmission is still needed^52^.

On the ecological side, consensus models (like the simple majority-rule model presented here) may be the first step towards decomposing suitability into something more epidemiologically-relevant. Development of alternative consensus models should aim to further clarify the level of suitability beyond the simple binomial categorization offered by ENM methods alone. The inclusion of a temporal component (i.e., the use of the conservative Mordecai projections of suitability for mosquitoes) enables some decomposition of the ENM results. The Mordecai *et al*. model illustrates that transmission is unlikely to be a year-round property of most areas, especially in temperate zones, and our exercise shows that reducing the months of possible transmission does significantly reduce total outbreak size. Time-specific ecological niche models have been used with great success to predict the dynamics of dengue^53^, another *Aedes*-borne disease, and have been applied as a proxy for Zika risk^26^. However, these models will need to be developed specifically for Zika as more data become available, and time-specific ecological niche models will pose an additional challenge for consensus building with mechanistic time-sensitive models like Mordecai *et al*.'s. Finally, we suggest the frameworks underlying consensus models should be adaptable as additional occurrence data is made available. ENMs are typically presented as static instantiations of dynamic processes, whether they describe species ranges of the transmission niches of emerging infectious diseases. The ability of these models to contribute to our understanding of pathogens entering novel regions or hosts will hinge upon their flexibility in incorporating near-real-time data^19^. The computational frameworks for dynamic, updating niche models exist^54^, but are an unexplored frontier in eco-epidemiology.

## Methods

### Ecological Models

Three studies have been published using ecological niche models (ENMs) to map the possible distribution of Zika virus, using a different combination of occurrence data, environmental predictors, and statistical approaches^12, 16, 17^. Their models suggest varying degrees of severity, especially as measured within the United States (Table 1). Other models have also been widely used in epidemiological work as a proxy for the distribution of Zika, such as an ecological niche model of *Aedes aegypti* and *Ae. albopictus*^29^ (fairly commonly used, e.g.,^51, 55^; or see^56^, which presents its own *Aedes* ENM that becomes a risk map of Zika transmission), or dengue-specific niche models (recently used by Bogoch *et al*. in two separate publications^26, 57^). Most ecological niche models indicate the range of Zika virus should be more restricted than that of its vectors, and published evidence suggests there may be significant differences between the known and potential distributions of dengue and Zika^12^, so we exclude these proxy methods from our study and focus instead on modeling studies that explicitly use Zika occurrence data.

#### Carlson et al

Carlson *et al*.^12^ developed an ensemble niche model constructed using the R package BIOMOD2. The resulting model uses seven of ten possible methods (general linear models, general additive models, classification tree analysis, flexible discriminant analysis, multiple adaptive regression splines, random forests, and boosted regression trees), notably omitting maximum entropy (MaxEnt). Their primary model uses only occurrence data from outside the Americas, but here we adapt their secondary model which incorporates data from Messina *et al*. (below) to show the lack of the sensitivity of the method to that additional data, especially in the United States. The only environmental predictors used are the BIOCLIM dataset^58^ and a vegetation index (NDVI). The final model threshold was selected to maximize the true skill statistic, with a selected value of 0.271 used in the original study to produce a binary suitability map. In the Carlson *et al*. model, suitable range for Zika virus is predicted to be limited to the southern tip of Florida and small patches of Los Angeles and the San Francisco Bay area. Only a total of 13 counties have any suitable area in this model; at the county scale, this model has the greatest concordance with observed outbreak patterns of autochthonous transmission in the United States during 2016.

#### Messina et al

Messina *et al*.^16^ use an ensemble boosted regression trees approach with a global dataset of occurrence points primarily from South American outbreak data. The model incorporates prior information about *Aedes* distributions. For example, pseudoabsences are preferentially generated in areas of lower *Aedes* suitability. Their model uses six environmental predictors: two direct climate variables, two indices of dengue transmission based on temperature (one for *Ae. albopictus* and one for *Ae. aegypti*), a vegetation index (EVI), and a binary land cover classifier (urban or rural). Messina *et al*. select a threshold of 0.397 that marks 90% of occurrence data as suitable (10% omission). Their model predicts that suitable range for Zika virus encompasses a substantial portion of the Gulf Coast, including the entirety of Florida and as far west as eastern Texas. Their study is also the first to estimate population-at-risk, placing the global figure at 2.17 billion people.

#### Samy et al

Samy *et al*.^17^ use MaxEnt to build four sub-models with different combinations of environmental predictors. The first is a conventional ENM approach using environmental predictors (precipitation, temperature, EVI, soil water stress, “aridity,” and elevation). The second, a more unconventional approach in the niche modeling literature, separates out socioeconomic predictors (among them population density, night light from satellite imagery, and a function of expected travel time called “accessibility”). The third uses all the same as the first model but with three added layers (land cover and suitability for *Ae. aegypti* and *Ae. albopictus*); finally, in model 4, all variables are included and we use that here as the representative case of the alternative Samy formulations. (In an additional sub-analysis, we compare these four models and show the impacts of these variable selection choices on downstream epidemiological forecasts.) For all, the model threshold is selected based on a maximum 5% omission rate for presence data, and also projects high environmental suitability in the Gulf region, very similar to that of Messina *et al*. This model also produced isolated suitable patches based on social factors, which predominantly occur at urban centers.

#### Mordecai et al

Mordecai *et al*.^10^ produced a Bayesian model of transmission of *Aedes*-borne viruses (dengue, chikungunya, and Zika) in the Americas that we adapt as a mechanistic geographic forecast for subsequent analyses. In their main model, an *R*_0_ modeling framework is constructed based on models for vector borne diseases, building upon the Kermack-McKendrick *R*_0_ model for malaria^59^. In this model, the majority of parameters describing the life cycle of mosquitoes and parasite development within the mosquitoes are sensitive to temperature. Mordecai *et al*. used data derived from the literature to parameterize the shape of the temperature response for each temperature sensitive parameter. These are based on laboratory observations of *Aedes aegypti* and *Aedes albopictus*, and infections with dengue, chikungunya, and Zika at constant temperatures through the range of possible values. Because these are bioenergetic functions, curve fitting exercises to derive appropriate models of the non-linear relationships underlie the parameterization of the overall transmission model. A non-linear overall relationship between transmission (*R*_0_) and temperature is fitted in a Bayesian inference framework, and from it two endpoints of a “suitable range” can be extrapolated within which *R*_0_ > 0. Those ranges can be adjusted for different levels of posterior probability, and can be used as a suitability threshold that can be projected onto gridded temperature data, producing binary monthly maps of suitability (which can be aggregated to year-round possible presence). In the Mordecai *et al*. publication, the most conservative probability level (> 97:5%) was then mapped onto long-term mean monthly average temperatures in the Americas, derived from Worldclim data^58^, to estimate the number of months transmission was possible for *Ae. aegypti* and *Ae. albopictus*^10^. Additional maps were also constructed of the number of months of possible transmission for *R*_0_ > 0 at posterior probabilities of 50% and 2.5%, and are found in the supplemental material. Here, we use all three probability levels from the *Ae. aegypti* model, to project the terms of the number of months of predicted transmission potential by mapping the model onto WorldClim temperature gridded data for long-term monthly minimum and maximum temperatures (six possible combinations).

#### Consensus Mapping Methods

In a preliminary effort to present a consensus forecast based on current ecological understanding, we use two alternative methods to develop county-scale predictions from the models included in our analysis. The first (“majority rule”) excludes the Mordecai model, and simply applies a majority rule to the binary thresholded Carlson, Messina, and Samy county shapefiles (i.e., any county with agreement between a majority of the niche models for either presence or absence). In the second model (“seasonal majority rule”), we take the counties predicted by the majority rule method and restrict their suitability to the months predicted in the strictest Mordecai model (97.5% confidence) for minimum temperatures. That process excludes 13 of the counties deemed suitable according to the simple majority rule, but which are predicted to be unsuitable year-round in the Mordecai model.

### Epidemiological Model

To simulate potential Zika outbreaks in the United States, we adopt the modeling framework used by Gao *et al*., which incorporates both sexual and vector-borne transmission^60^. We selected Gao *et al*.'s framework because, while fairly simple, it includes a number of important features of the epidemiology of Zika, including the high rate of asymptomatic cases, and lingering (primarily sexual) transmission by post-symptom “convalescent” cases. Because the transmission term is normalized by dividing by total population size, the model itself is scale-free. Thus, the values associated with each compartment could be represented as proportions rather than the number of individuals. The model divides the human population into six compartments with levels: susceptible (*S*), exposed (*E*), symptomatically (*I*_1_) or asymptomatically (*A*) infected, convalescent (*I*_2_), and recovered (*R*), where he *h* and *v* refer to the human host and mosquito vector populations, respectively:

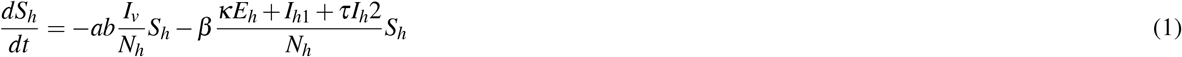

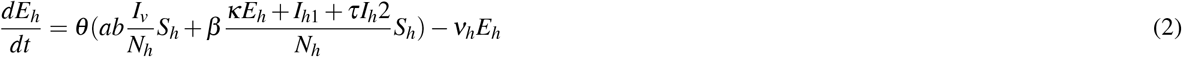

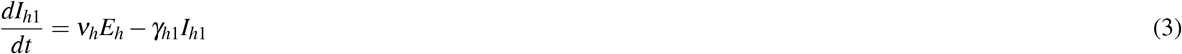

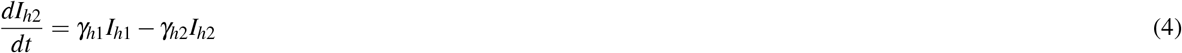

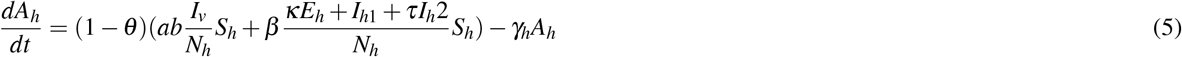

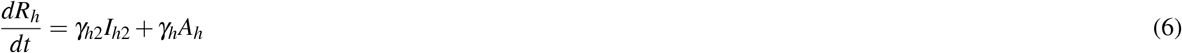

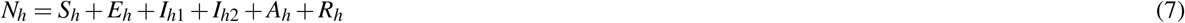

Vectors are governed by a complimentary set of equations but only divided into susceptible, exposed, and infected classes:

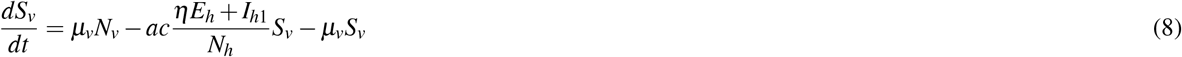

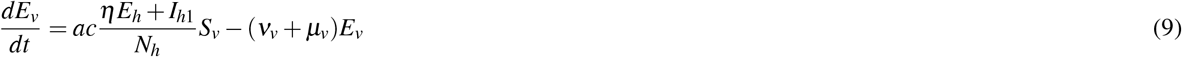

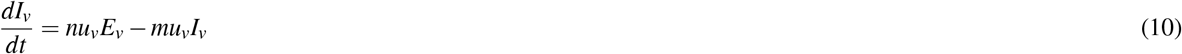

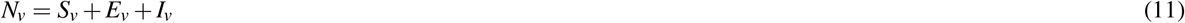

Rather than use the fitted parameters from any given country's outbreak, parameters for the above models were randomly generated from a set of uniform prior distributions specified by Gao *et al*. as reasonable priors based on the literature (Table 2). Evidently, these models are significantly discrepant with outbreaks in the continental U.S. so far, with fewer than 300 cases of local transmission recorded in 2016 (and in fact, our simulations are far more severe in terms of final case burden than estimates for Brazil or Colombia). However, the purpose of applying this epidemiological model across the spatial extent predicted by each niche model is both to illustrate the uncertainty that goes unstated in presenting such ENMs and to intimate the necessity of developing and parameterizing these models in concert.

### County-Level Simulations

In our main models, every spatial projection of Zika risk was summarized at the U.S. county scale, such that if a single pixel within a county polygon was projected to be suitable under a given model, the county was marked suitable for outbreaks. This assumption clearly overestimates population at risk, but environmental suitability is often aggregated to the county scale in order to develop Zika models for the U.S.^19, 51^. For the Mordecai models, the maximum value (months suitable per year) of all pixels within a county was assigned as the value. For example, if a single 25 km^2^ cell in a particular county was suitable for a single month, simulations were run for one month with mosquitoes present and the remaining 11 with a mosquito population of zero. While this approach has the potential to overestimate populations vulnerable to mosquito-borne transmission, it adds a number of key strengths. Aggregating information at the county scale absorbs some of the relative spatial uncertainty of predictions at the pixel scale, and may account for source-sink dynamics for vector-borne outbreaks driven by heterogeneity in vector density and competence. Moreover, the county level is one of the finest scales at which public health infrastructure is likely to decide whether interventions like vector control are necessary. Finally, sexual transmission can spread from cells with suitable vectors to vector-free areas, and as a function of both sexual transmission and underlying mobility, outbreaks are therefore unlikely to be contained to a given pixel. Previous work has similarly used the county scale to study risk factors and model outbreak risk for Zika^19, 51^, and we follow their precedent.

Population data for each county was taken from projections to the year 2016 based on the 2010 United States Census, and were set as the total susceptible human population at the start of a year. The mosquito population was set at five times the baseline human population, the middle of the range selected by Gao *et al*.^60^. While other studies have used a lower ratio^20^, we set mosquito populations (the only parameter we explicitly fixed from the Gao model) as high as we did because many simulations with lower mosquito populations faded out immediately, and setting a higher ratio made the impacts of model differences more immediately apparent. Outbreaks were simulated stochastically at the county level using the Gao. model, initiated with a single infected person per county. We randomly selected a value for each of the parameters in the Gao et al. model for which a range was provided, using a uniform distribution (Table S1). For each modeling model, 100 simulations were run in each county designated as suitable. For the three ENMs, county models were run for a “model year” (twelve months of thirty days each), and had no interactive effect on each other. For the Mordecai models, the full vector- and sexually-transmitted epidemic models were run for the number of months (thirty days each) that were predicted suitable. After that period, the total vector population *N_v_* was set to 0, effectively ending vector-borne transmission, but models continued so that sexual transmission was ongoing up to 360 days. All simulations were run in R 3.3.2, and all scripts and county simulation data are available as supplementary files.

#### Within-County Heterogeneity

In a final set of analyses, we examine the impact of how risk is aggregated at the county scale. Fine-scale population data does exist for the world from multiple sources^61^, but at the resolution niche models are often generated, clear problems exist. Running models on a pixel-by-pixel basis would likely be computationally prohibitive in many cases (including this one); moreover, in the context of sexual transmission, models that do not explicitly include human movement between nearby pixels might produce results that make little sense. While vector movement may be fairly minimal, human movement likely produces mixing at broader geographic scales for both vector-borne and sexual transmission. Aggregating niche models to a county-level suitability is one solution to the problem, and has the added benefit of plausibly absorbing some of the uncertainty among different ENMs. However, this also has the clear tendency to overestimate population at risk; to examine how strongly this affects models, we include an additional set for Carlson, Messina, and Samy (model 4) where susceptible population is scaled down linearly by the proportion of the county marked suitable in each model. This, in itself, adds another layer of neutral assumptions (populations are treated as having a uniform distribution within counties) but might also produce less drastic differences between outbreak trajectories. The results of that analysis are given in Table 3 and supplemental Figures S1, S2, and S3.

## Data Availability Statement

Data generated or analyzed during this study are made available on Figshare: doi.org/10.6084/m9.figshare.5514961.v1

## Acknowledgements

We thank Lewis Bartlett, Eva Harris, and others for helpful comments and guidance. This work was supported by the Rutgers University Center for Discrete Mathematics & Theoretical Computational Science (DIMACS) Mathematics of Planet Earth 2013+ Workshop on Zika & DIMACS MPE 2013+ Workshop on Appropriate Complexity Modeling of the Impacts of Global Change on Ecosystems, and the Centers for Disease Control and Prevention Epidemic Predictions Initiative (CDC EPI), a Center funded by NSF (EF-0553768). SJR was also supported by NSF DEB EEID 1518681, NSF DEB RAPID 1641145, and CDC grant 1U01CK000510-01: Southeastern Regional Center of Excellence in Vector-Borne Diseases: the Gateway Program. This publication was supported by the Cooperative Agreement Number above from the Centers for Disease Control and Prevention. Its contents are solely the responsibility of the authors and do not necessarily represent the official views of the Centers for Disease Control and Prevention.

## Author contributions statement

CJC conceived of the study, and contributed the ecological niche models. SJR contributed the mechanistic models. ERD designed code for epidemiological simulations. CJC and ERD contributed to figures. All authors contributed to the writing and editing of the manuscript.

